# KAGE 2: Fast and accurate genotyping of structural variation using pangenomes

**DOI:** 10.1101/2023.12.23.572333

**Authors:** Ivar Grytten, Knut Dagestad Rand, Geir Kjetil Sandve

## Abstract

Structural variation is known to play an important and often overlooked role in regulating disease and traits, but accurately detecting structural variants from sequencing data has traditionally been difficult. However, recent improvements in high-quality genome assembly along with methodological advancements in pangenome creation have opened up the landscape for methods that use such pangenomes for structural variant calling and genotyping. We here present KAGE2, which accurately and efficiently genotypes structural variation by exploiting the availability of pangenomes that represent known variation in a population. Through comprehensive benchmarking, we highlight limitations of existing methodology and show that KAGE2 is more accurate and considerably faster than existing methods.

## Introduction

While the study of genetic variation traditionally has been focused on SNPs and short indels, we now know that structural variation plays a much more important role than previously assumed [1,2,3]. Thus, being able to cost-effectively detect structural variation in a genomic sample is of great importance. Traditionally, detecting structural variation has been done by mapping short-read sequences to a reference genome, which yields much lower accuracy than equivalent methods for SNPs/short indels, since reads originating from the structural variants are less likely to map to the reference [4]. A solution has been to use longer reads [5], and while there exist several tools [6,7] that are able to accurately detect structural variation based on long reads, it comes with a markedly increased sequencing cost.

A more promising “hybrid” approach, leveraging both the accuracy of long-read methods and the cost-effectiveness of short-read sequencing, has recently gained traction [9]. The idea of this hybrid approach is to create a high-quality database (referred to as a pangenome) of some select genomes and their variants using accurate long-read sequencing. With such a pangenome [10], variants in an individual of interest can be characterized by genotyping variants in the pangenome, i.e. for every known variant in the pangenome, estimating the individual’s genotype for that variant. This can be done with short-read sequencing by comparing sequenced reads directly to the pangenome (e.g. through graph-based mapping-techniques [11,12]). This approach allows for discovering any variation in a sequenced individual that is already represented in the pangenome, and circumvents the issue of mapping short reads to a linear reference genome, which also is known to lead to reference bias [13]. Another benefit of this approach is that known population structure encoded in the pangenome can be used to improve accuracy, e.g. by “imputing” variants that are difficult to call using information from other variants [9,14].

This genotyping approach has previously shown promising results for SNPs and short indels [9,14,15,16], but its use has been limited for structural variation, since few good pangenomes with structural variation have been available. However, the recent release of a high-quality human pangenome containing 47 haplotype-resolved individuals [8] now make this approach much more relevant for structural variant genotyping of humans (and there are also similar ongoing projects for other species [17]). As part of this human pangenome release, the authors showed that the genotyper PanGenie [9] is able to genotype structural variation of a new individual (an individual not present in the pangenome) with fairly high accuracy. A problem with PanGenie is, however, that its runtime scales quadratically with pangenome size (number of individuals in the pangenome). This is a problem as pangenomes are expected to grow significantly in size in the years to come.

We here present KAGE2, which expands recent methodology for SNP/indel genotyping to allow pangenome-based genotyping of structural variants. We here show that KAGE2 is more accurate than existing methods for genotyping structural variation in addition to being considerably faster.

## Results

We present KAGE 2, which extends our previously published genotyper KAGE to enable genotyping of structural variants. KAGE2 genotypes structural variants through a strategy similar to SNP/indel genotyping, by exploiting a pangenome that represents known structural variation in a population. The variants present in the population are represented by kmers, allowing KAGE2 to infer structural variant genotypes by comparing these variant kmers to kmers present in the reads for the individual that is being genotyped. We refer to [14] for an overview of the original KAGE genotyper. The main methodological changes from KAGE to KAGE2 are how kmers are selected from the pangenome in order to represent structural variants (see Methods). KAGE2 also implements direct support for imputation using GLIMPSE [18] (see Methods).

In the following, we present various benchmarks that compare the accuracy of KAGE2 against the existing genotypers PanGenie and Bayestyper, using the recently published high quality draft human pangenome [8]. We do this using a leave-one-out setup where an individual is removed from the pangenome and genotyped using the remaining pangenome (see Methods).

### KAGE is faster and more accurate than existing methods

We run KAGE2 and previously proposed genotypers with various read coverages (average number of reads per base pair), and compute accuracy as F1 score using Truvari [19]. As can bee seen from Figure 1a, KAGE2 outperforms other methods for all read coverages. We also ran the same experiments with 5 other randomly picked individuals, which all showed similar results (see Supplementary Figure 1).

**Figure 1:**
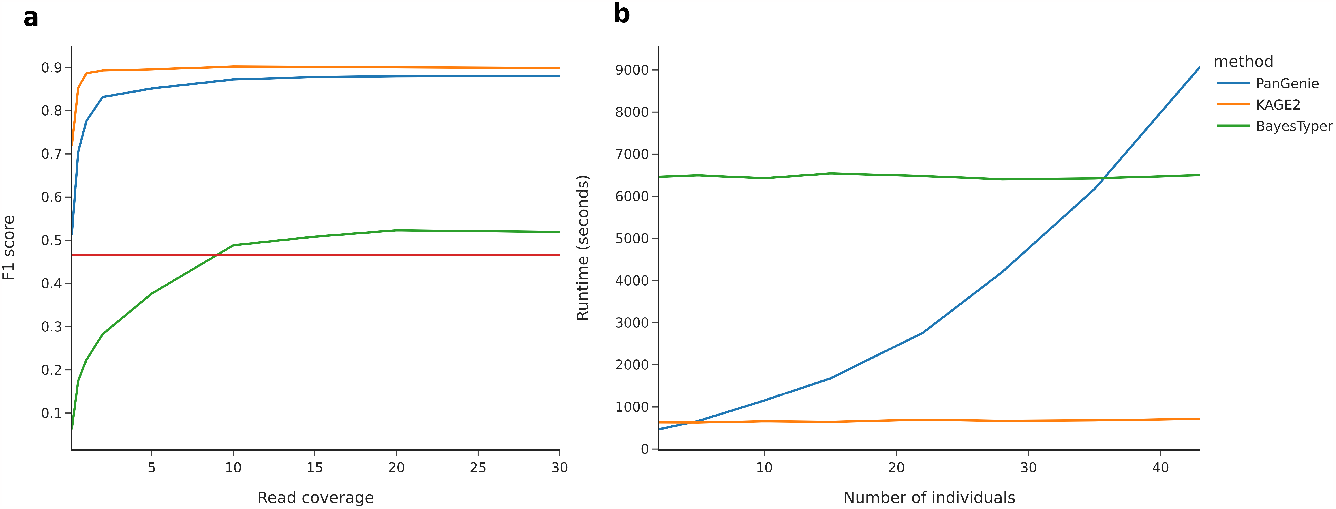
a) Accuracy on genotyping structural variants when removing a random individual from the pangenome and genotyping that individual. Baseline (guessing) is a naive baseline genotyper that does not use any reads, but instead “guesses” the genotype just by using population priors (Methods). b) Runtime as a function of pangenome size (number of individuals in the pangenome) when running on chromosome 1 of the Draft Human Pangenome [8]

Figure 1b shows the runtime of the genotypers as a function of pangenome size (number of individuals in the pangenome, see Methods). KAGE is considerably faster than competing methods, and its runtime is independent of pangenome size. Note that Bayestyper’s runtime is the same for all pangenomes, which is because it does not use any information from the individuals in the pangenome. Since PanGenie uses a Hidden Markov model with possible haplotype path as states, its runtime increases drastically with the number of individuals. The time spent for creating indexes for PanGenie and KAGE is not included in the runtime (but listed in the Supplementary Material).

### KAGE performs well on different variant types/regions

To investigate the observed performance in more detail, Figure 2 shows the genotyping accuracy stratified by type of genomic regions and type of variant (using regions defined by GIAB [20]). KAGE2 is generally the best-performing genotyper across all variant-types and regions.

**Figure 2:**
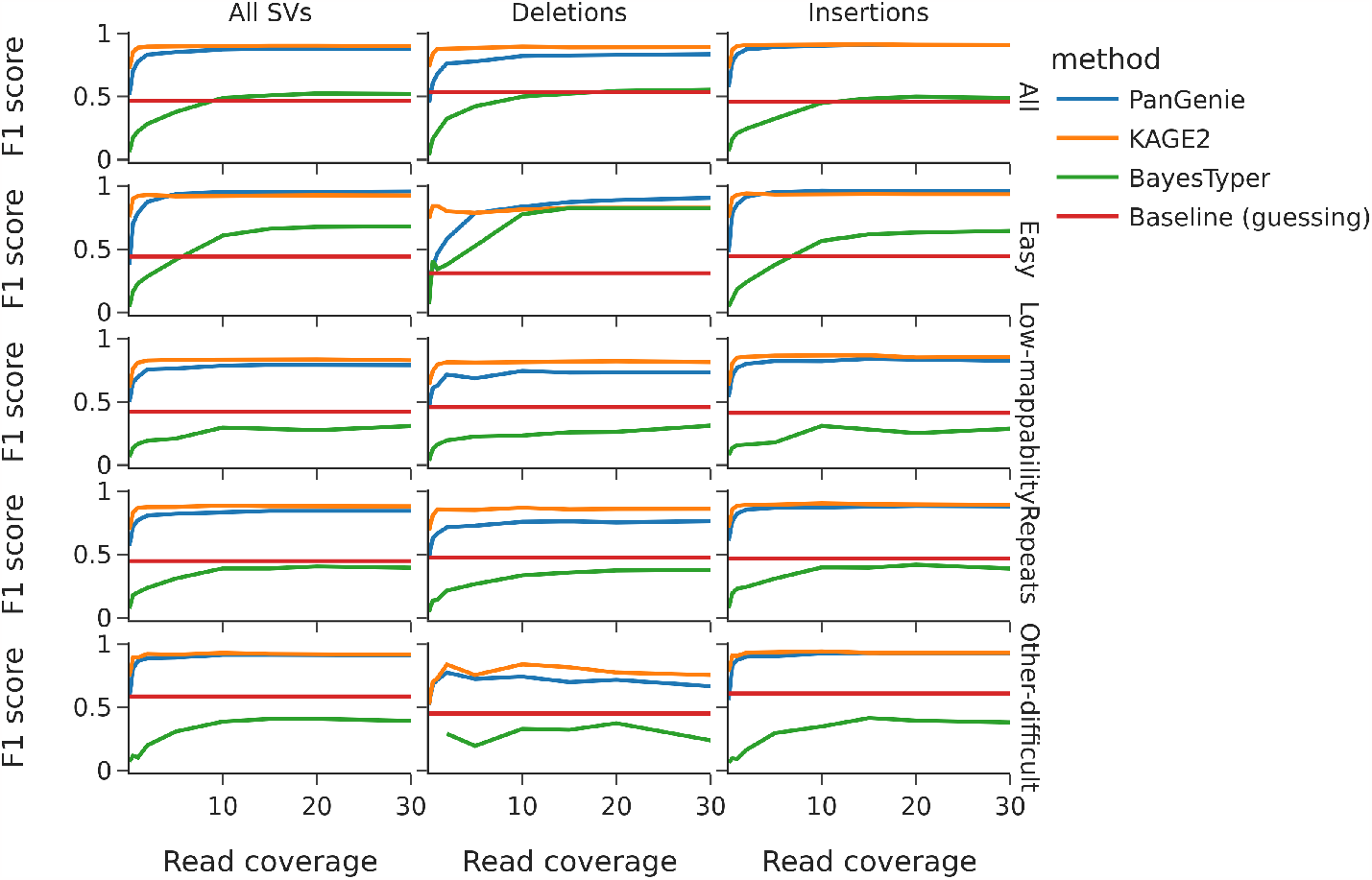
Accuracy for deletions and insertions in various types of genomic regions.

### Pangenome size matters

A limitation of pangenomic genotypers as opposed to de novo variant callers is that the genotypers can only detect a variant if it is also present in the pangenome used for genotyping. Thus, larger pangenomes (created from more individuals) opens for higher recall, though with a risk of lower precision. Figure 3 shows accuracy as a function of the number of individuals in the pangenome. As can be seen, accuracy increases as more individuals are added, although with a diminishing return for every additional individual. This indicates that it is likely worth the effort to invest in creating even larger pangenomes, and that having methods like KAGE that have runtime independent of pangenome size is important. Interestingly, the baseline genotyper performs worse when the pangenome grows, which is likely because the inclusion of more rare variants increases the number of false positive calls.

**Figure 3:**
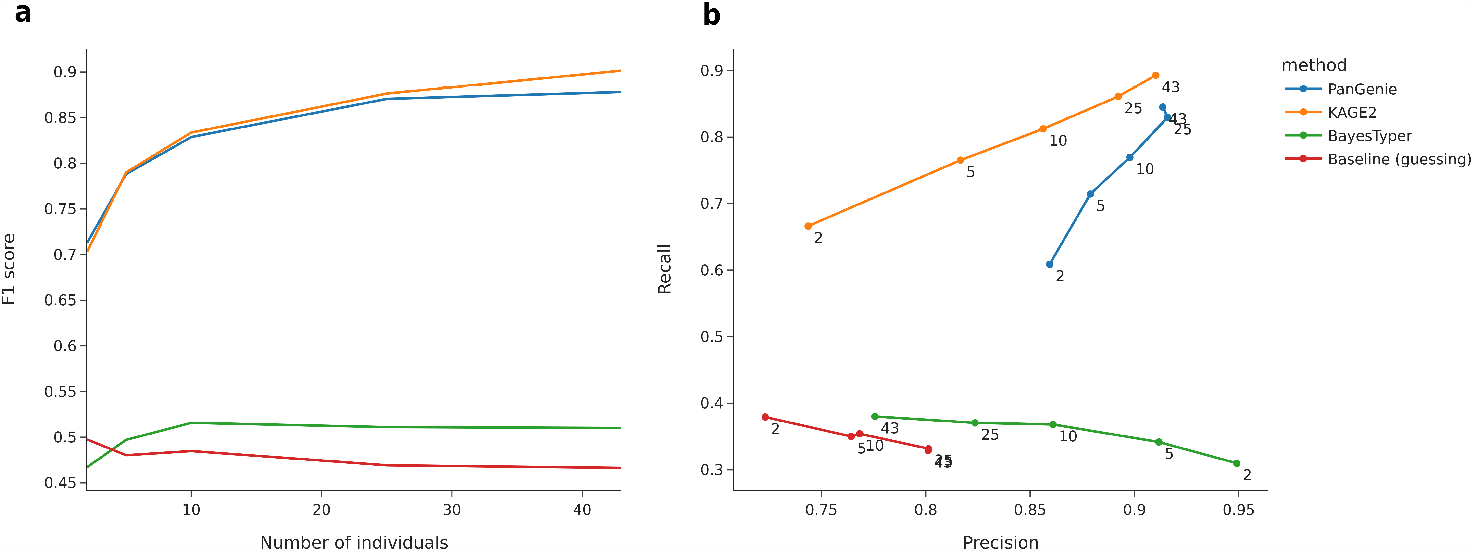
Accuracy as a function of pangenome size (number of individuals in the pangenome) on all structural variants.

### SNPs/indels help calling structural variants

We also investigate whether the genotyping of structural variants is improved by including more SNPs and indels in the pangenome. We do this by creating a variety of pangenomes, where we for each pangenome filter away SNPs/indels that have allele frequency lower than a given threshold. As can be seen from Figure 4, both PanGenie and KAGE benefit from more SNPs/indels.

**Figure 4:**
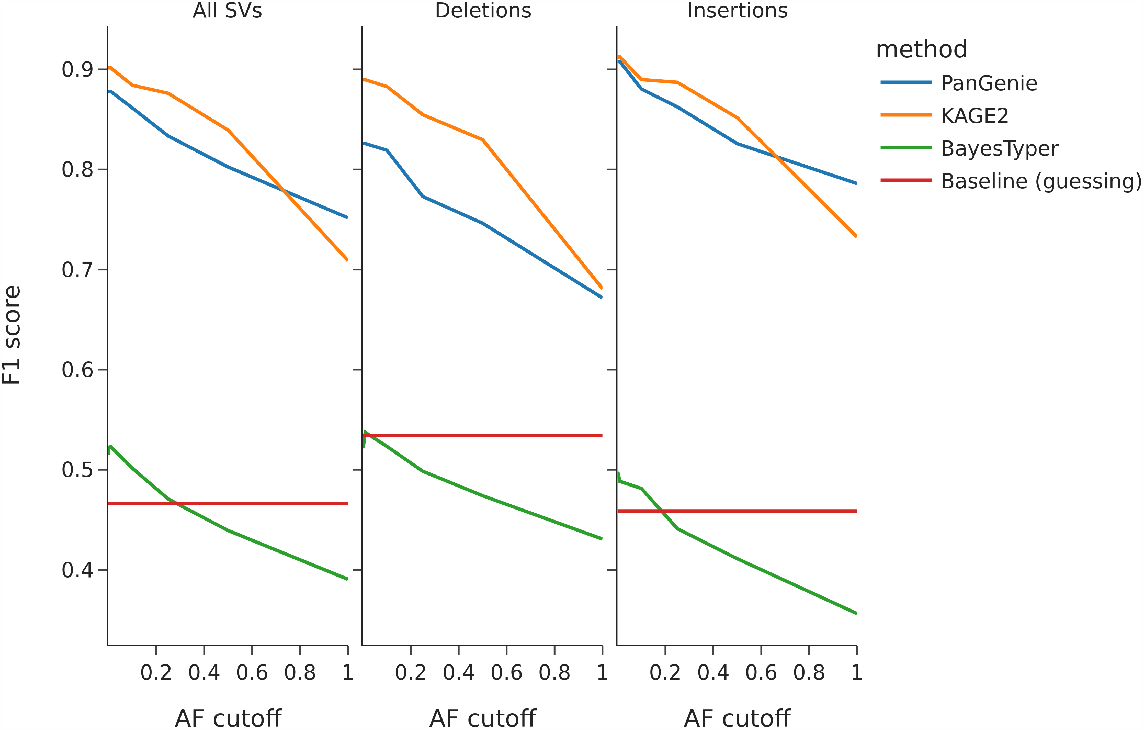
Accuracy when SNPs/indels have been filtered using various allele frequency cutoffs. All SNPs/indels with allele frequency lower than the cutoff have been removed, meaning that the number of SNPs/indels decreases as the cutoff increases. The baseline genotyper has constant accuracy, since it only genotypes variants using population priors (which for structural variants do not change when SNPs/indels are removed).

## Discussion

We have presented KAGE2, a genotyper that is able to efficiently and accurately genotype structural variation from short reads by using a pangenome representation of a population. We find it useful to view the problem of calling structural variants based on pangenomes as consisting of of two sub-problems: 1) The problem of extracting useful information from reads (for KAGE and PanGenie this is based on kmers), and 2) the problem of using information about individuals represented by the pangenome to improve prediction (imputation). While PanGenie solves both these problems jointly using a Hidden Markov model, KAGE2 approaches these modularly as two separate problems. This allows us to use external software (GLIMPSE) for the imputation model. A benefit of defining these as two problems is that it allows for the design of a modular solution where existing tools can be used. If another better imputation tool than GLIMPSE would be made available at some point, which is a likely development, the imputation part can be easily exchanged within KAGE2.

A surprising result is the high level of accuracy achieved by KAGE2 (with use of GLIMPSE imputation) when read coverage is low (0.5-2x) (Figure 1). In fact, there is almost no gain in accuracy when read coverage is increased beyond 5x (towards the more commonly used 30x). We speculate that the reason is that most structural variants are difficult to call based on read information alone, and that accurate genotyping is mainly driven by imputation. The fact that accuracy goes drastically down when there are fewer SNPs/indels in the pangenome (Figure 3) further shows how important it is to include SNPs/indels in the pangenome.

While the recently released draft human pangenome only consists of 47 individuals, the Human Pangenome Reference Consortium has announced plans to release a pangenome with 350 individuals in 2024 [8]. As it is clear that pangenomes will only continue to grow in the future, we believe KAGE2 to be an important contribution to the field not only because of its current improvement over state-of-the-art methods, but also because it will be able to computationally scale and exploit the increased information expected from future, larger pangenomes.

## Methods

We here describe how KAGE2 is implemented and how the experiments were performed. When referring to structural variants in this manuscript, we lean on the commonly used definition of structural variants being variants where either allele contains 50 or more bases.

### KAGE2 implementation

The main difference between KAGE and KAGE2 is that KAGE2 employs an improved strategy for picking kmers to represent variants, which is needed since structural variants are often multiallelic and contain repetitive sequence. Through experimentation, we found it important to select kmers that have low frequency locally (within the variant) as well as globally (in the pangenome). Kmers are chosen by looking at multiallelic variant sites, so that all overlapping variants in a region are grouped and looked at together as one multiallelic variant. For each such multiallelic site, we first find all kmers covering every allele of this multiallelic site. Among these kmers, the aim is to find a kmer to represent each allele. For each kmer candidate we assign a score that is based on how often this kmer is observed in the pangenome globally, as well as how often it is observed locally at the multiallelic variant. We minimize the global score, but found it more important to minimize the local score, as this is important to be able to separate the alleles of a multiallelic variant when genotyping. Thus, we weigh the local score twice as much as the global score, and for each allele pick the kmer with lowest score. For some variant sites, several alleles will end up being represented by the same kmer (because no unique kmers can be found). In such cases, KAGE may still be able to draw useful information from the reads, because the expected frequency of each kmer in input reads may be different depending on which combination of alleles the individual has. Selecting and scoring kmers is done efficiently using BioNumPy [21] and NumPy [22].

### Using GLIMPSE for imputation

A central part of how the original KAGE genotyper performed genotyping was the use of estimated genotype likelihoods between preselected pairs of variants to guide genotyping, which used information from the population about how genotypes of pairs of variants are “correlated”. This is a simple form of imputation [23], and while the original KAGE genotyper worked well for SNPs and indels, we have found this approach to be less successful for structural variation. Instead, we have found GLIMPSE [18] to work well for imputing structural variants and thus use GLIMPSE as default imputation model. Whenever we are referring to KAGE2 in the results, we refer to KAGE2 with imputation done by GLIMPSE. The implementation of KAGE2 also still supports genotyping using the previous builtin imputation model for cases where this would be desired.

### Benchmarking

For all results presented in the manuscript, KAGE2 has been using the human draft pangenome [8]. This pangenome is built from whole-genome assembly of 47 individuals and is expected to represent most of the individuals’ SNPs, indels and structural variants with high accuracy. We perform experiments by removing one individual from the pangenome (i.e. remove all information stemming from the given individual) and genotype that individual using the remaining pangenome. This mimicks a real-case scenario, where we have an existing pangenome and are to genotype a new sample that is not present in the available pangenome based on short-read sequencing for the new sample. In this way, we can compare the genotypes we predict to the original genotypes (those annotated in the original human draft pangenome based on expensive long-read sequencing) to get an estimate of genotyping accuracy of KAGE2 based on short-read sequencing.

In the experiments presented in this manuscript, we only run the methods on one human chromosome with simulated reads. This allows us to explore several parameters efficiently. In the Supplementary material, we run a subset of the experiments on the whole human genome with reads from the 1000 Genomes Project [24], showing that results on the whole genome with non-simulated reads give similar results.

We chose to mainly compare KAGE2 to PanGenie, as PanGenie has previously shown that it is both faster and more accurate when genotyping structural variants than other methods. Also from our own experience, PanGenie is the only existing available method that is able to genotype structural variants using pangenomes in reasonable time with decent accuracy. We also include BayesTyper [25] as a reference, which is a method that works somewhat similarly to KAGE2 and PanGenie by representing variants with kmers and counting those kmers in the reads.

We have implemented a comprehensive pipeline for benchmarking structural variant genotyping accuracy, which allows us to explore how a variety of parameters (such as read coverage, allele frequency, number of individuals in the pangenome, read error rate) affect accuracy. Our pipeline can be found at https://github.com/bioinf-benchmarking/sv-genotyping. Experiments with various parameter configurations can be easily configured using this pipeline.

### Measuring genotype accuracy

Measuring the accuracy of structural variant genotyping is less trivial than with SNPs and short indels, as identical or near-identical structural variants can be represented in multiple ways. In previous benchmarks [8,9], accuracy has been defined using weighted genotype concordance, which briefly explained is an average of the precision (correctly assigned genotypes divided by the total number of variants assigned that genotype by the genotyper) for each of the three possible genotypes (0/0, 0/1 and 1/1) (see [9] for formal definition). We find two issues with this metric:

1. Equal weight is put on all genotypes (0/0, 0/1 and 1/1). Since the number of variants in each group is skewed (there are many more 0/0 than 0/1 and 1/1), this metric is easily boosted by assigning 0/0 to most of the variants, and only calling 0/1 or 1/1 for variants that are predicted with high certainty. This allows to achieve a very high precision on 0/1 and 1/1 without sacrificing much precision on 0/0, meaning that the weighted average will get high although the overall accuracy is low (a large number of false 0/0 calls). We believe it is better to use the more standard way of measuring accuracy by using recall and precision, as is also recommended by others [26].
2. The weighted genotype concordance requires exact genotype match to mark a genotype call as correct. In practice, most structural variants in the draft human pangenome are multiallelic (sometimes with as many alleles as the number of haplotypes in the pangenome), and many alleles are almost identical within a multiallelic variant. Thus, requiring the genotyper to find the exact allele among many almost identical alleles is in our opinion too strict, and in practical settings what we are interested in is whether we are calling an allele that is similar enough (within some threshold) to the truth. The tool Truvari implements a way to measure accuracy with some slack on allele matching, and we refer to [19] for a further discussion on this. We have thus chosen to use Truvari when measuring genotyping accuracy instead of the matching criterias used in [8] and [9].

While previous benchmarks [8,9] ignore genotype errors that are due to variants of the individual being genotyped not being present in the pangenome, we instead count these as errors (false negatives). This is because we want the experiment to be as close to a real-world scenario as possible, where we genotype an individual and are interested in how many variants we find (recall) and the accuracy of those calls (precision). A pangenome that lacks variants present in the individual will lead to lower recall since fewer variants can be detected by the genotyper, and we want the results to reflect this limitation.

### Baseline genotyper

In the experiments we have included a “baseline” genotyper. This genotyper is implemented by running GLIMPSE with uniform genotype likelihoods as input, meaning that they are a result of GLIMPSE only using population priors to do genotyping and no information from sequenced reads.

## Code and data availability

KAGE2 is available at https://github.com/kage-genotyper/kage/. Instructions for how to reproduce the experiments provided here are available at https://github.com/bioinf-benchmarking/sv-genotyping/kage-experiments.md. The pangenome used in all experiments was downloaded from https://zenodo.org/records/6797328.

