## Supplementary Material for "KAGE 2: Fast and accurate genotyping of structural variation using pangenomes"

### Supplementary Figure 1

Figure 2 rerun with 5 random individuals (random seed in benchmarking pipeline ranging from 1-5).

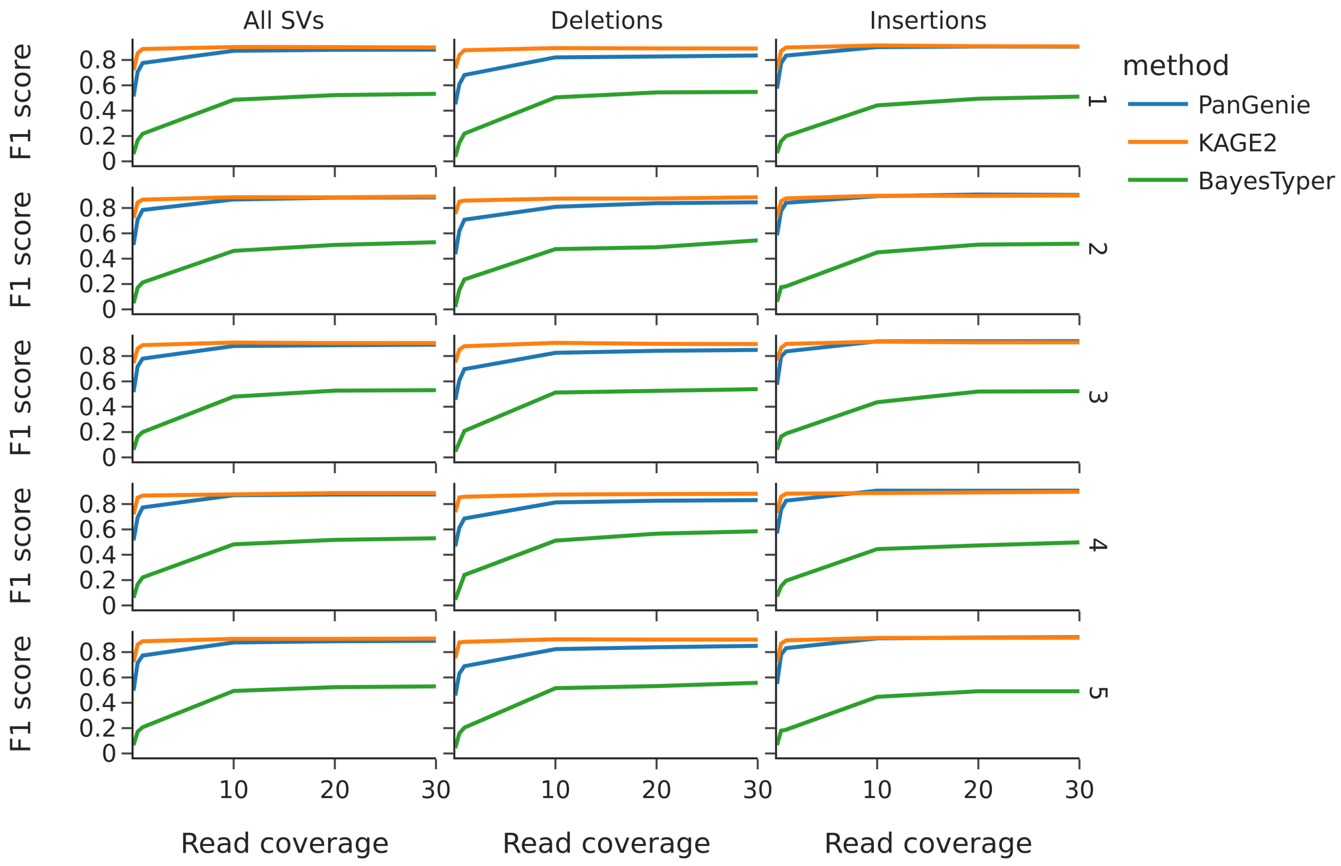


### Supplementary Figure 2

Figure 2 rerun with reads from 1000 Genomes Project on the whole human genome for one individual.

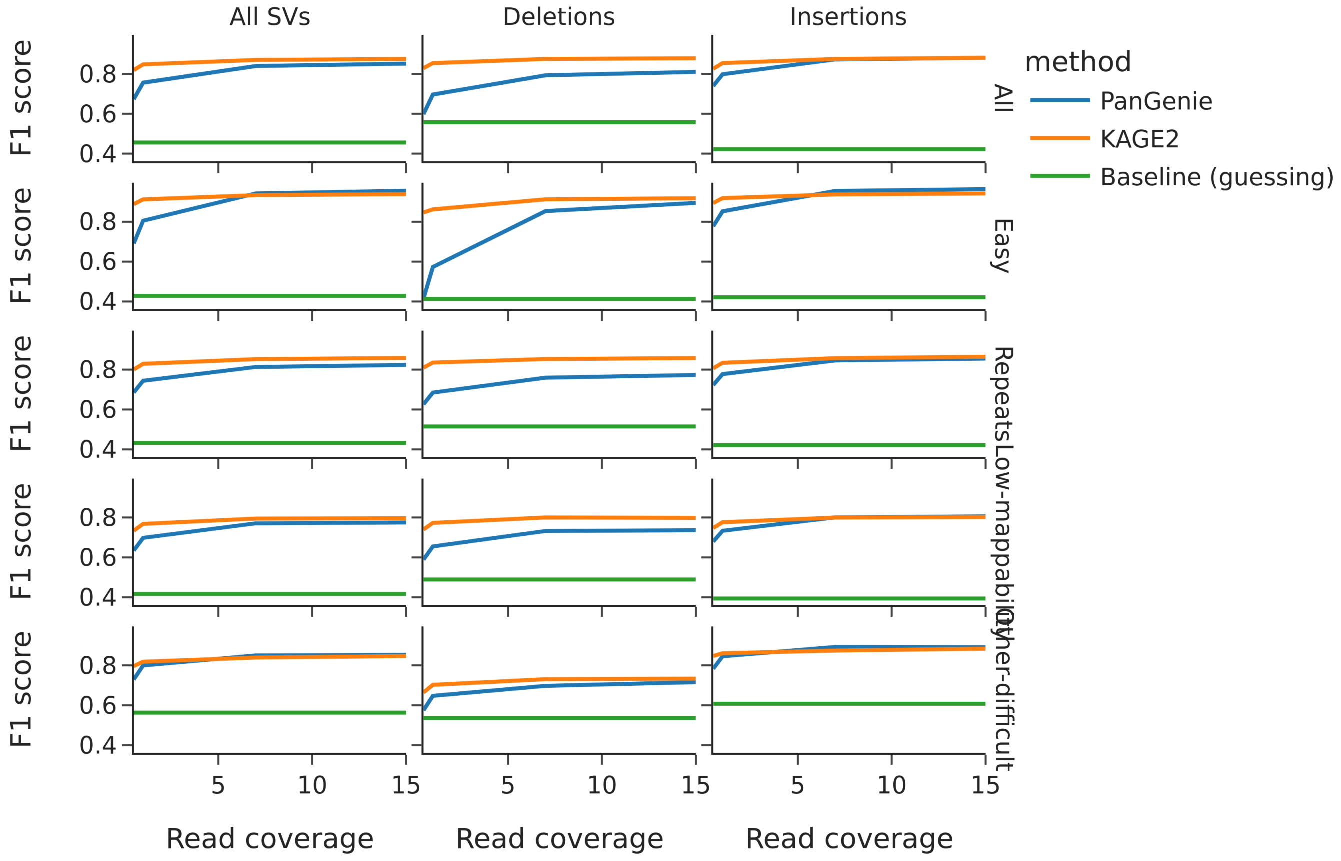


### Indexing time

The following table shows the time spent creating indexes for the whole Draft Human Pangenome.

| | **Time spent (h:m:s)** |
|---|---|
| KAGE | 23:22:48 |
| PanGenie | 2:17:49 |
